# Weber’s law and natural inference

**DOI:** 10.1101/2024.08.10.607448

**Authors:** Jeffrey Beck, Ingmar Kanitscheider, Guillaume Dehaene, Alexandre Pouget

**Affiliations:** Department of Neurobiology. Duke University, NC; Department of Basic Neuroscience, University of Geneva, Switzerland; Center of Learning and Memory and Department of Neuroscience, The University of Texas at Austin, Austin, TX; Department of Brain and Cognitive Sciences. University of Rochester. Rochester, NY; Gatsby Computational Neuroscience Unit. London. United Kingdom

## Abstract

For most extensive sensory variables such as speed or numerosity, the discrimination thresholds of human subjects are proportional to the value around which the discrimination is performed, a scaling known as Weber’s law. Many theories have been proposed for this law, which all rely on the assumption that neurons are noisy. By contrast, we argue here that noisy neurons are not required to explain Weber’s law. Instead, we propose that it is the unavoidable consequence of the statistics of natural sensory inputs. In natural environments, sensory measurements are typically scaled by global variables such as contrast in vision or loudness in audition. These global scaling parameters induce positive correlations among measurements which in turn lead to Weber’s scaling. This theory makes testable experimental predictions and accounts for the fact that tuning curves to speed and numerosity in vivo are approximately log normal.

## Introduction

The difference in numerosity that human subjects can reliably discriminate between two sets of items is proportional to the average numerosity in the display (Fig. 1a). Thus, if a subject can detect a difference of 3 items when the average number of items in the display is 10, the discrimination threshold would be close to 6 for displays with about 20 items. This is known as Weber’s law, a law that has been observed for a wide variety of extensive variables (variable for which addition makes sense) such as line length, surface, speed of motion, elapsed time, distance traveled and weight comparison (Consweet and Pinsker, 1965; Fechner, 1966; Gaydos, 1958; Indow and Stevens, 1966; Oberlin, 1936). If we denote the discrimination threshold *δs* and the stimulus around which the discrimination is being performed, *s*, then Weber’s law implies that the ratio *δs* / *s* (also called the Weber fraction) is a constant function of *s*. If the ratio *δs* / *s* is constant, then psychometric curves are scale invariant: they depend on ratios, not absolute values (Fig. 1b-c).

**Figure 1:**
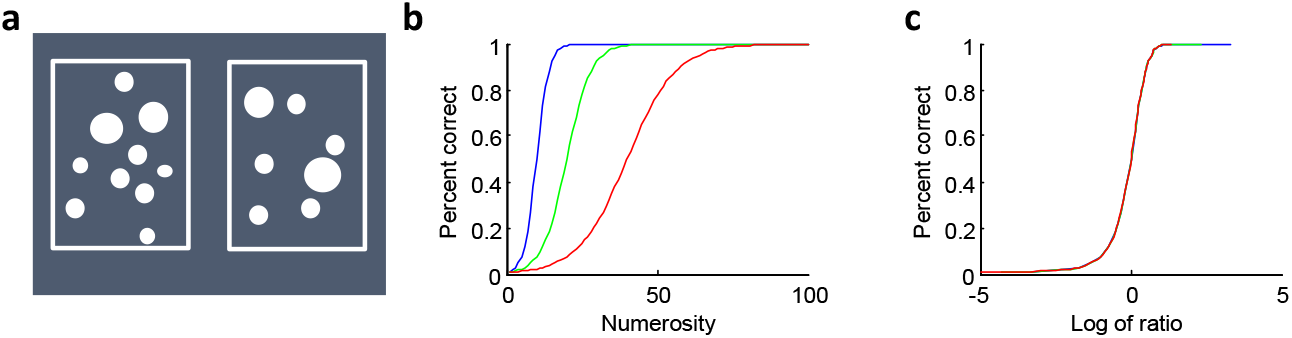
Weber’s law. **a**. Typical display for a numerosity discrimination task. Subjects have to judge whether there are more dots on the left than on the right. **b**. Hypothetical psychometric curves following Weber’s law for numerosity discrimination. The curves show percentage of correct responses as a function of the numerosity for discrimination around a reference value of 10 (blue), 20 (green) and 40 (red). Note that the slope of the psychometric curve become shallower as the reference value increases. **c**. When the psychometric curves shown in panel **b** are replotted as a function of the logarithm of the ratio of the numerosity over the reference numerosity (10, 20 and 40), the curves are superimposed. This is a consequence of Weber’s law: performance depends only on ratios of values as opposed to absolute values.

A common explanation for Weber’s law relies on the idea that neurons respond proportionally to the logarithm of *s*, or have Gaussian tuning curves to the logarithm of *s*, and are corrupted by noise with variance independent of *s* (Dehaene, 2003; Dehaene and Changeux, 1993; Nover et al., 2005). It is easy to show that the maximum likelihood estimate of *s* obtained from such a neural code follows a Gaussian distribution across trials with mean equal to *s* and a standard deviation proportional to *s* (in the limit of large numbers of neurons). Therefore, the discrimination threshold based on this maximum likelihood estimate would also be proportional to *s*, because discrimination thresholds are proportional to standard deviations. This in turn implies that the ratio *δs* / *s* would be constant, as required by Weber’s law. There are several variations of this idea (e.g. Deco et al., 2007) all of which hinge on one critical assumption: neurons are noisy.

Others have suggested that Weber’s law reflects the scale invariant nature of many natural variables (Chater and Brown, 1999). For instance, the power spectrum of natural images is scale invariant: it stays the same regardless of the magnification factor applied to an image. According to these theories, Weber’s law is the consequence of encoding optimally scale invariant variables with noisy neurons. In the same vein, it has been proposed that Weber’s law is due to a rescaling of the input to optimize coding given the limited coding capacity of neurons (Gallistel, 2011; Gallistel, 1990). These explanations also rely on the assumption that neurons are noisy since the noise is what sets the limits on the coding capacity of individual neurons.

The noisy neuron assumptions is problematic because it is far from clear that the noise corrupting neuronal responses has a major impact on behavioral variability and therefore on discrimination thresholds (Beck et al., 2012; Moreno-Bote et al., In press). Here, we explore a different explanation, one that does not rely on nonlinear processing by noisy neurons but on the presence of global nuisance parameters that induce positive correlations in sensory measurements. According to this theory, Weber’s law would still be observed even if neurons were noiseless because Weber’s law is a consequence of the statistics of natural inputs. To simplify our presentation, we first focus on the perception of numerosity, before discussing a more general theory, along with experimental predictions.

### Weber’s law for numerosity

Consider the problem of counting *N* white spots shown against a black background (Fig 2). We assume that all spots have the same size with luminance *x*_*i*_ with mean *μ* and standard deviation *σ* (Fig 2, left column). An estimate of numerosity can be obtained by taking the integral of the luminosity across the display:

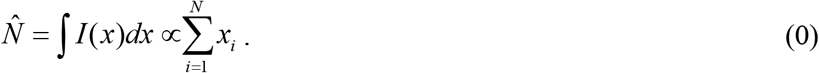

**Figure 2:**
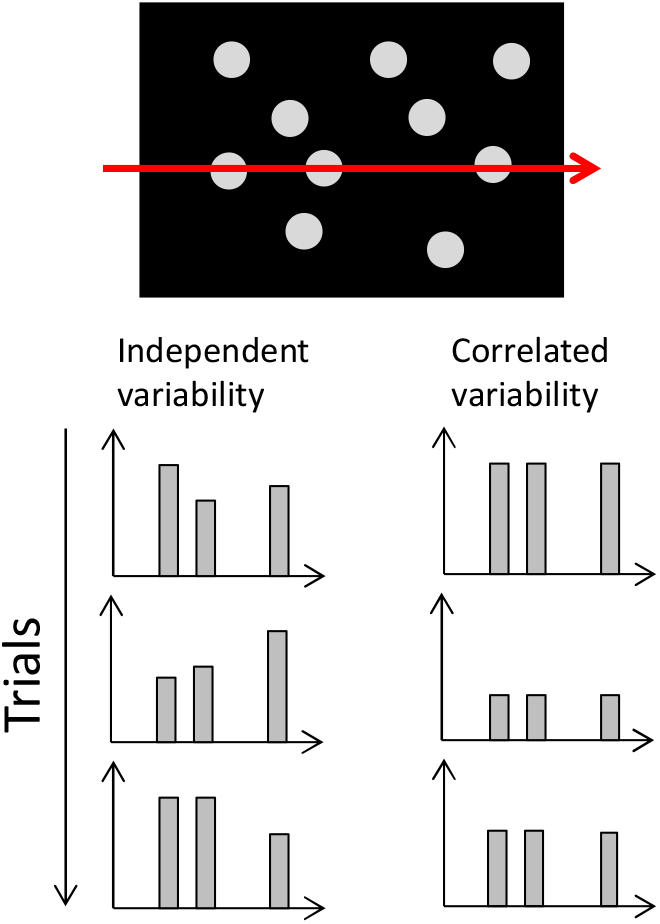
Independent versus correlated variability. The bottom plots show examples of luminance profiles across trials taken along the red arrow on the visual display on top. Left column: examples of luminance profiles corrupted by independent zero mean gaussian noise. Right column: same as on the left but with fully correlated noise. Note that in this case, all bars go up and down together. This kind of variability leads to Weber’s scaling.

Such an estimate, however, does not follow Weber’s law. To see this, we need to compute the ratio of the standard deviation over the mean of the estimate. According to Weber’s law, this ratio should be constant. The mean of 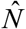 is proportional to the number 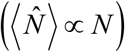 of dots but the standard deviation is proportional to 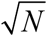. Indeed, we are simply adding *N* independent random variables, in which case the variance of the sum is equal to the sum of the individual variances which implies that the variance is proportional to *N*, or equivalently, that the standard deviation is proportional to 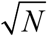. Therefore the ratio of the standard deviation over the mean is not constant but inversely proportional to 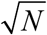.

The simple reasoning however is only valid if the overall contrast is the same in all images. If this is not the case, the luminance of each dot is given by *y*_*i*_ = *gx*_*i*_, where *g* is a random variable corresponding to global contrast which varies between images. This model of natural images is called a scale mixture model, and is known to capture some fundamental properties of natural statistics(Schwartz and Simoncelli, 2001)). Because of the shared contrast random variable, *g*, the measurements *y*_*i*_ are now positively correlated (Fig. 2, right column). As a result, the variance of the sum of the *y*_*i*_ is no longer simply proportional to *N*. Instead, it follows:

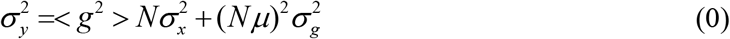

where <*g*> and 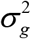 are the mean and variance of the contrast. For large *N*, the variance scales with the number of items squared, which corresponds to Weber’s law. By contrast, for small *N*, the variance is proportional to *N* as before, which predicts that Weber’s law should break down for small numbers. This is indeed what has been found experimentally (Cordes et al., 2007).

This shows that the presence of global nuisance variables, such as contrast, is sufficient to account for Weber’s law. However, estimating numerosity from total luminance is clearly problematic when contrast varies from trial to trial, as is the case in natural images. Indeed, the integral of the luminance is not simply proportional to numerosity but to the product of numerosity and contrast 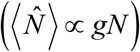. This problem can be alleviated by normalizing this sum by an estimate of the global contrast obtained from the image, which we denote *ĝ*. The estimate of numerosity would now be given by:

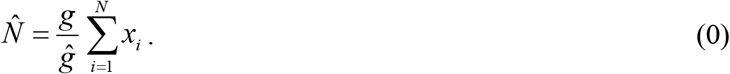

This normalized estimate of numerosity still follows Weber’s law because all the variables *x*_*i*_ are scaled by a common nuisance variable *g* / *ĝ* (Fig. 2a), which induces the positive correlations that led to Eq. (0). However, dividing the estimate of numerosity with an estimate of contrast as we do in Eq. (0) is not necessarily the optimal solution to normalize out contrast. A better approach consists in marginalizing out contrast. As we show in section 1 of the Supplementary Material, under reasonable assumptions for the distribution of g, this also results in an estimate of numerosity that follows Weber’s scaling for large numerosity.

One might object that Weber’s law is observed even when the contrast of the dots is maintained from trial to trial, that is, even when *g* is not a random variable. Under such conditions, if we assume that subjects are aware that the contrast is fixed, then the ratio *g* / *ĝ* should not vary across trials, in which case Weber’s law should no longer hold. This is indeed correct but this requires that subjects know that the contrast is constant in this specific experiment. This will likely require extensive training on the subject’s part since contrast constantly varies from one image to the next in natural environments. Therefore it is quite likely that neural circuits automatically estimate contrast on every trial. Estimating contrast however is difficult. For one thing, there is no accepted universal definition of contrast that applies across all images; instead, the definition of contrast tends to be stimulus specific (Reinagel and Zador, 1999). Moreover, it is not easy to determine where to estimate contrast and over which area. This problem is even worse when taking into account the fact that the positions of the objects can change on each trial, as well as the position of the eyes and the position of the attentional spotlight. If the estimate of contrast is based on the part of the image that is foveated or attended, this image will vary from trial to trial even if the image is the same. Solving this difficult inference problem implies that the estimate of contrast, and therefore the ratio *g* / *ĝ* in Eq. (0), will vary from trial to trial even when g is constant in the image, thus leading to Weber’s law. This is also true if the subject uses the full Bayesian model for inference (section 1 of the SI).

Note that there are other global nuisance parameters for numerosity, beside contrast and eye position, that require some form of normalization. For instance, even when counting something as simple as circles on a screen, it is important to normalize the total surface area by an estimate of the per item surface area or to normalize total perimeter squared by total surface area, to ensure that the counts do not double when the size of the objects double. This implies that there are multiple multiplicative nuisance parameters that could induce positive correlations among measurements, all of which contribute to the Weber fraction.

### General theory

This theory of Weber’s law can be readily generalized to other variables beside numerosity. For visual variables in which the front-end computation involves a linear combination of pixel values, such as speed (Adelson and Bergen, 1985) or line length estimation, contrast acts as a global nuisance parameters for the same reasons as the ones we discussed for numerosity (and so does surface area). Indeed, the scaling of the variance for positively correlated measurements (Eq. (0)) is not limited to straight sums but generalizes to any positive linear combinations of measurements. As such this theory applies to the estimation of any extensive quantity (extensive quantities are quantities for which addition makes senses, such as weight or length).

For example, for weight comparison (Weber, 1978), Weber’s law might be due to the co-contraction of muscles that can act as a global scaling signal. Indeed, the arm keeps the same posture when the agonist and antagonist muscles are co-contracted by the same factor. This co-contraction factor induces positive correlations among the proprioceptive signals or the efferent copies that are presumably used to estimate the weight. For distance traveled, if the estimation is based on the number of steps (as is the case for ants, for instance,(Wehner and Srinivasan, 1981)), any variation in the mean distance of a step, because of muscle fatigue, or change in the terrain, will also correlate the step counts.

Moreover, learning could also induce positive correlations. For instance, consider a task in which subjects are shown a display with two sets of intertwined dogs and cats, and are asked to determine whether there are more dogs than cats. In this case, each image must first be classified as either cat or a dog before it can be added to the counters. If the subject uses a linear classifier, the classification would operate on a decision variable of the form **w**·**v**_*i*_, where **v**_*i*_ is a vector of sensory measurements for object *i*, and **w** is a vector corresponding to the linear classifier. As learning proceeds the subject adjusts the weight vector after each trial to optimize performance. There is already experimental evidence that such online learning takes place on a trial by trial basis (O Donchin et al., 2003). The updated weight vector can then be used to process the items in the next display. Since all objects are classified with the same weight vector on each trial, the variability in **w** from trial to trial induces positive noise correlations among the object classifications, thus leading once again to Weber’s law (see section 2 of the Supplementary Material for details).

This last factor might play a particularly important role in Weber’s law for time estimation. Weber’s law in time estimation might be due to the fact that subjects count the ticks of an internal clock (Matell and Meck, 2000) whose rate vary from trial to trial, while being fairly constant during a single trial (Treisman, 1963). The rate fluctuations would act like the contrast factor in our numerosity example, correlating positively the pulses of the clock, and leading to Weber’s law. But why would the rate vary from trial to trial? One possibility is that subjects constantly calibrate their clocks both with one another and with clock like signals from the external world (Ahrens and Sahani, 2011). A clock is indeed useless if not properly calibrated. A sensible way to calibrate the clock would be to make small adjustments after each use of the clock, or each trial in an experiment.

### Experimental predictions

If Weber’s law is due to the presence of global nuisance parameters, one might expect that increasing the variability of these nuisance parameters should increase the Weber fraction. Thus, the Weber fraction might increase if the brightness of the dots in an image varies from trial to trial. This intuition, however, is not necessarily correct. Recall that if the subjects normalize for the contrast (Eq. (0)), the Weber fraction will now depend on the variance of the ratio *g* / *ĝ*. If the variability in *g* also follows Weber’s law (as is the case for instance for luminance), this ratio has a fixed variance regardless of the variance in *g*. This would predict that increasing the variability in contrast should have little impact on the Weber’s fraction, which is indeed what has been found experimentally (Burgess and Barlow, 1983).

This theory also predicts that the response of integrators in the brain should generally exhibit Weber’s scaling (standard deviation proportional to *N*) unless all nuisance parameters are properly controlled, in which case it should follow a distribution in which the standard deviation proportional to 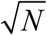. Moreover, in the latter case, Weber’s law should reemerge if the experimenter introduces a global nuisance parameters in the display.

Since there are many potential nuisance parameters, controlling these parameters will undoubtedly require extensive training. For instance, to get rid of the learning effect, the weight vector must stop fluctuating. This could happen if subjects learn that the category boundary (the weight **w** in the previous section) does not fluctuate over time. Since this rarely happens in real life, the amount of data required to conclude with high certainty that the classification boundary is stationary is necessarily exceptionally large.

However, one result appears to indicate that it is indeed possible to achieve this goal. This result comes from the analysis of the variance of the rate of LIP neurons involved in the temporal integration of sensory evidence for decision making. As we pointed out, for the simple integrator receiving independent evidence over time, the variance of the integrator should scale with its mean (see paragraph below Eq. 1), which is what we call counting noise and is sub-Weber in its scaling. For a temporal integrator, this implies that the variance should grow linearly with time, which is indeed the case for LIP neurons in highly trained monkeys (Churchland et al., 2011). In this particular task, the animals were trained to discriminate between rightward versus leftward motion which is a very natural classification boundary to learn. It is therefore quite possible that the weight vector used by the monkeys to distinguish between these two directions does not fluctuate after enough training. We predict however, that the variance of the LIP neuron will show Weber scaling when the animal are trained to determine whether the direction of motion is clockwise or counterclockwise from say 37 degrees, or when they are trained to determine whether the coherence in the display is more or less than 10% to the right. In both cases, learning such an arbitrary boundary is difficult, and there is evidence that animal keeps adjusting their classification boundaries for such difficult tasks, even after extensive training (Mendonca et al., 2012).

### Implications for Neural Coding

One theory of Weber’s law is based on the idea that neurons receive inputs that are not subject to Weber’s law but encode these inputs with log normal tuning curves corrupted by internal noise (Dehaene, 2003; Nover et al., 2005). This is indeed consistent with the observation that neurons in vivo have, to a first approximation, log normal tuning curves, that is, the width of the tuning curves to the log of the stimulus is roughly constant (Nieder et al., 2002; Nover et al., 2005).

In contrast, we are suggesting that sensory inputs are already subject to Weber’s law. This theory does not rely on the existence of log normal tuning curves, which raises the question as to why such tuning curves are found in vivo. In this section, we show that these log tuning curves might in fact provide an optimal code for sensory inputs following Weber’s law. In other words, log normal tuning curves might be the consequence of the scale nuisance parameters that lead to Weber’s law rather than the cause of Weber’s law itself.

We derive this result by seeking a population code that maximizes the logarithm of the Fisher information averaged over the prior distribution of the stimulus. Fisher information is a natural measure to use because the inverse of Fisher information provides a lower bound on the discrimination thresholds of an ideal observer of the neural activity (Papoulis, 1991). In addition, the log of Fisher information can be related to Shannon information for Gaussian distributed posterior distributions (Brunel and Nadal, 1998). Following Ganguli and Simoncelli (2014), we perform the maximization under two constraints: 1-that there is a finite number of spikes available for coding, and 2-a finite number of neurons. Moreover, we assume that the spike counts of individual neurons conditioned on their inputs are drawn on every trial from an independent Poisson distribution and that the sensory inputs entering this network follows Weber’s law.

Under such assumptions, we found that the code is optimized when the tuning curves to the log of the stimulus have a constant width (Fig 3a, see Section 3 of the supplementary information for details), as has been reported in vivo. When plotted as a function of the stimulus (as opposed to its log), these tuning curves have a width roughly proportional to the preferred stimulus of the neurons (Fig 3b). Importantly, this result is independent of the choice of the prior distribution of the sensory inputs. The prior distribution only influences the density of the tuning curves. For instance, for a prior proportional to 1/*s*, the optimal density is proportional to 1/*s* (Fig 3a-b). Importantly, the Weber fraction of the optimal code is also approximately constant over the range covered by the tuning curves (Fig. 3c). As a result, the code follows Weber’s law just like its input.

**Figure 3:**
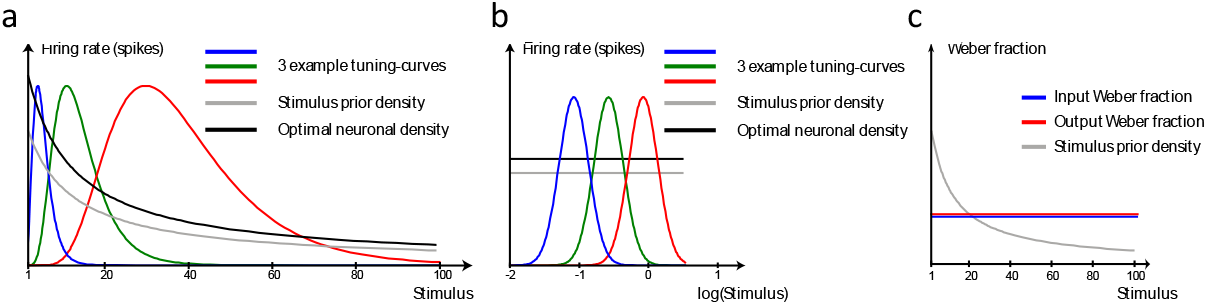
Optimal neural code for sensory inputs following Weber’s law. **a**. Three examples of optimal tuning curves for encoding a sensory input following Weber’s law. The widths of the tuning curves increase with their peak value. The solid black line show the optimal density for the stimulus prior distribution indicated by the grey line. In this example, the prior distribution is *s*^−1^, as in the main text. **b**. Same tuning curves as in **a** but plotted as a function of the log of the stimulus. The width of the tuning curves is constant, indicating that the optimal tuning curves are log normal as has been reported in vivo for variables like speed and numerosity. Log normal tuning curves might therefore be a consequence of optimally encoding inputs that follow Weber’s law. **c**. Weber fraction for the input (blue) and the output (red) of the neural code as a function of the stimulus. The Weber fraction is a constant function of the stimulus by construction. The Weber fraction is also constant for the output thus reflecting the fact that the output of the optimal neural code also follows Weber’s law. With sufficient number of neurons and evoked spikes, the output Weber fraction is almost identical to the input fraction. The grey line still shows the prior distribution of the stimulus used in the simulations.

Intuitively, this result is easy to understand. In order to minimize information loss, the network should devote more coding resources to inputs that are more precise and more likely. The optimal population reflects both of those influences. Since the input follows Weber’s law, its precision decreases with its mean value. Correspondingly, the precision of the optimal code drops off for higher values of the encoded stimulus. This drop is caused both by the decrease in neuronal density for higher values and the increase in the width of the tuning curve to the stimulus.

Note also that while noise did not play a role in our theory so far, this particular result is only valid if individual neurons are noisy and their noise, conditioned on the input, follows an independent Poisson distribution. For a very different noise structure (such as Gaussian additive noise with fixed variance), the optimal code might not involve log normal tuning-curves.

## Discussion

We have proposed that Weber’s law is due to the presence of global scaling variables either in natural stimuli (e.g contrast) or in the data generated by biological hardware (e.g. change in the mean size of steps due to muscle fatigue). The presence of such global scaling parameter creates positive correlations in measurements which then lead to Weber’s law for a large class of linear or nonlinear functions of the measurements. We are not the first one to recognize that positive correlations in measurements lead to Weber’s law. For instance, Treisman (1964) made a similar proposal but, interestingly, he suggested that the positive correlations were a property of neuronal noise, while we argue that is a property of the data received by the nervous systems. In the time domain, several authors (Ahrens and Sahani, 2011; Gibbon et al., 1984; Treisman, 1963) have also proposed that a rate varying clock would lead to time estimates following Weber scaling but to our knowledge, they never suggested that this could constitute a general theory of Weber’s law, nor did they speculate on the origin of these fluctuations (such as a calibration of the clock as we suggest here).

The proposal outlined in this work stands in contrast with several theories of Weber’s law which rely on the combination of nonlinear processing (typically a log nonlinearity) followed by the addition of neuronal noise (Dehaene, 2003; Nover et al., 2005). The idea that neuronal noise (noise internally generated by cortical neurons) is the main limiting factor on human performance in psychophysics tasks is indeed popular but the evidence is not as strong as typically believed (Beck et al., 2012). Indeed, if neurons were stochastic devices and the stochastic fluctuations were independent, then these fluctuations could be easily averaged out. Correlations could change this picture, but there is currently no evidence that neural circuits create correlations that limits the information available in neural codes (they are claims that neural activity exhibit correlations that limits information (Zohary et al., 1994), but this does not mean that the correlations are generated internally (Moreno-Bote et al., In press)). In contrast, we have recently proposed that behavioral variability is due instead to two factors: suboptimal inference and variability in sensory inputs (by which we mean the actual physical stimuli in the external world, or the sensors like photoreceptors) (Beck et al., 2012). This paper makes the additional suggestion that the variability in the sensory inputs is such that optimal, and near optimal, inference algorithms results in Weber scaling, whether or not downstream neurons are noisy.

Of course, it might be the case that neurons are indeed noisy, and that this noise also contributes to Weber’s law. Thus, it has been proposed that optimal coding theory predicts Weber’s law for particular combinations of noise, cost function and prior distribution over stimulus values (Ganguli and E.P., 2010). The basic idea behind such theories is that the number of neurons encoding a value should be proportional to the a priori probability of this stimulus value. Assuming neurons are noisy, this results in a code in which precision is higher for more likely values which in turn can lead to Weber’s law if small values are more likely than large ones. The weakness of such a theory is that they are typically not robust to deviations from the assumed cost functions and prior distributions. For instance, for a population of neurons with independent Poisson noise, a code that maximizes the expected inverse Fisher information will exhibit Weber’s law if the prior distribution is a polynomial function with an exponent of exactly –4 (Ganguli and Simoncelli, 2014). A different choice of cost function (e.g. Mutual information, which corresponds to the expected logarithm of Fisher information), or a different prior distribution (even if it’s still a polynomial function but with a different exponent) may yield to a different scaling. In our theory, Weber’s scaling does not depend on a particular choice of cost function or prior distribution since it depends solely on input statistics. Moreover, we showed that the log normal tuning curves that have been observed in vivo optimizes the efficiency of the neural codes when the input follows Weber’s law. This result suggests that these log normal tuning curves are the consequence of Weber’s law rather than their cause, in contrast to previous models.

The theory presented here also could explain the emergence of Weber’s law in a recent model of numerosity based on deep belief networks (Stoianov and Zorzi, 2012). The authors reported that when a deep belief network was trained on a set of images made of varying numbers of black shapes, it develops numerosity sensitive neurons. Moreover, when trained on a numerosity discrimination task, the performance of the network depends only on numerosity ratios as expected for Weber’s law. The authors of the study do not speculate on the reasons for this behavior. It could be due to a combination of the nonlinearity and variability of the units (deep belief networks rely on stochastic units). However, the authors also show that the network behavior is well captured by a simplified set of equations which involves a normalization for surface area. As we showed in Eq. (0), this would be sufficient to explain the emergence of Weber’s law in this model.

One appealing aspect of our theory is that it explains why Weber’s law is observed even in a domain in which there is little reason to have such a scaling if it could be avoided. For instance, any foraging animal has a strong incentive to know with the highest precision its position with respect to its lair or nest so as to minimize the energy that will be spent to return home. Passing the estimate of distance traveled through a log transform before it is corrupted by noise makes little sense in this case, as this would increase the variance of the distance estimate. Instead, the animal should use a linear path integrator (Gallistel, 1990), which would yield a variance proportional to the distance, as opposed to the distance squared. However, our theory argues that the animal has no choice: even if it uses a linear path integrator correlations induced by nuisance parameters such as step size or foot friction will automatically induce Weber scaling.

To summarize, we propose that Weber’s law is a consequence of the statistics of natural sensory inputs, as opposed to a consequence of internal noise. If we are right, computer software for variables like speed or numerosity will also exhibit Weber scaling when dealing with natural images even if they are noiseless and do not make use of compressing nonlinearities such as log.

## Acknowledgements

A.P. was supported by a grant from the Swiss National Foundation #31003A_143707 and from the National Science Foundation #NSF1109366.

## Supplementary Information

### 1) Weber’s law as a result of marginalizing contrast

In this section, we show that Weber’s law still arises when we marginalize over contrast (or scale parameter) rather than divide by a noisy estimate of the contrast (Eq. 3. main text). To demonstrate the generality of this result, we will show that it can be obtained when the underlying variable of interest is assumed to be either a continuous quantity that is either Gamma or Gaussian distributed, or a discrete count. The approach we will take is to first show that the marginal posterior over the quantity of interest (*s* for continuous variables or *N* for discrete counts) has a variance which is proportional to the square of the mean. This implies that, if the generative model is correct and the observer is performing optimal inference, then her behavioral estimates should also obey Weber’s Law. However, in experiments where the scale nuisance parameter is held fixed, this generative model is not correct. When this is the case, we show that estimates generated from the posterior mean still result in Weber’s Law like behavioral estimates due to fluctuations in the animals estimate of the scale nuisance parameter as in Eq. 3 of the main text.

*Case 1: Numerosity Estimates*

We assume that subjects sum, *s*, of the luminance of the objects, *x*_*i*_, present on the screen is confounded by a scale nuisance parameter *g*. Thus the measured quantity is total luminance given by

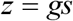

where

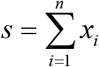

We assume that the underlying *x*_*i*_ are distributed according to:

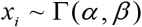

and our goal is to infer the number of objects *N* given the measurement *z*, that is, we want to infer the posterior distribution *p*(*N* | *z*) and then use the mean of this posterior distribution as the estimate of numerosity on each trial. We then show that the variance and mean of this estimate over trials follow Weber’s law.

**Figure 1:**
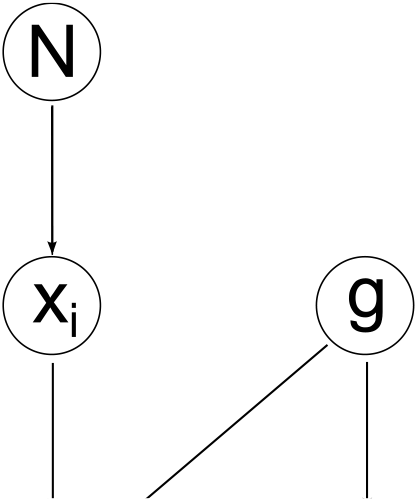
graphical model for numerosity.

Inferring *p*(*N* | *z*) is an underdetermined problem that can only be solved using additional information about the contrast *g*. We assume that the visual system is able to get this information from additionally observed image features which may include information obtained from past trials. This corresponds to performing inference on the graphical model in Fig SM1. We do not model these visual features explicitly but merely assume that these features specify a posterior distribution over *g, p*(*g* | *I*) which follows an inverse-Gamma function whose parameters depend on the current image, that is

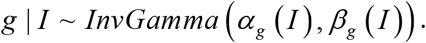

Assuming a flat prior on *N*, the posterior *p*(*N* | *z, I*) is proportional to the likelihood function *p*(*z* | *N, I*). The likelihood function can be obtained from *p*(*z, g, s* | *N, I*), by marginalizing out *g* and *s*, which leads to:

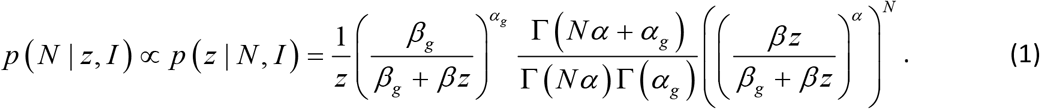

A posterior of this form is a one parameter generalization of the Negative Binomial distribution for *N*. In particular, when *α* = 1 it is easy to see that

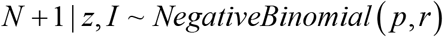

where

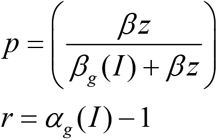

For negative Binomial distributions, the ratio of the standard deviation to the mean is simply 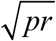 which is independent of z for large z. Therefore, when this generative model is correct, behavioral estimates of *N* obtained from the true posterior mean will exhibit Weber’s Law variability.

Weber’s law also persists in an experiment where this generative model is not correct, but is none-the-less being utilized by the subject to infer *N*. For instance, this situation arises in experiments in which the contrast is constant in all images, but the subjects assume that it varies.

First of all, we continue with the simple case *α* = 1. Let us assume that the subjects’ estimate on each trial, 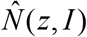, is given by the mean of the posterior distribution in Eq.(2):

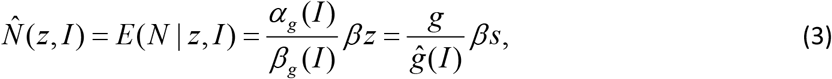

where *ĝ*(*I*) ≈ *β*_*g*_ (*I*) / *α*_*g*_ (*I*) approximately corresponds to the posterior mean estimate of contrast (when the image *I* is highly informative about the overall contrast of the image, both *α*_*g*_ (*I*) and *β*_*g*_ (*I*) will be large and *ĝ*(*I*) will be a good approximation to the posterior mean estimate of contrast.) The mean numerosity estimate is then given by

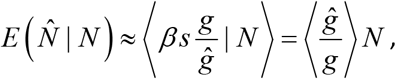

which indeed scales linearly with the true numerosity.

To determine whether this estimate follows Weber’s law, we compute the variance of the estimate over trials conditioned on the true numerosity *N*. Note that 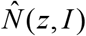 in Eq. (3) can be written as the product of the two independent random variables 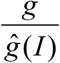 and *βs*. We therefore obtain

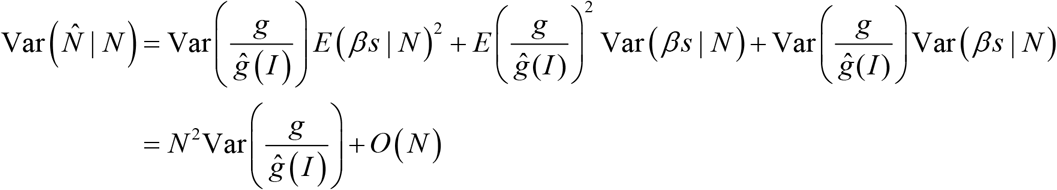

Therefore, for large *N*, this estimate follows Weber’s law, i.e., its variance is proportional to its mean squared, with a Weber fraction 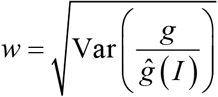.

When *α* is not equal to one it’s a bit more complicated, but the same conclusion can be drawn. To that end, assume that even though numerosity is discrete, in the limit of large *N*, we can replace the discrete posterior by a continuous distribution.

The posterior *p*(*N* | *z, I*) derived from Eq. (1) can be written in a closed form if *α*_*g*_ (*I*) is a positive integer. In the case *α*_*g*_ (*I*) = 1 it will be a Gamma distribution with parameters

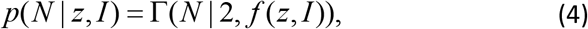

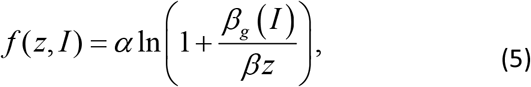

For *α*_*g*_ (*I*) = 2, 3,… the posterior will be given by a mixture of Gamma distributions with different shape but identical scale parameters,

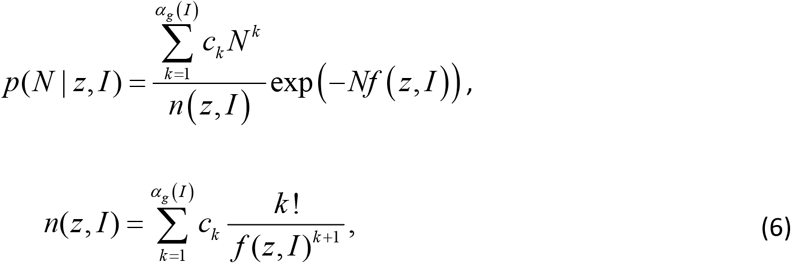

where *c*_*k*_ are the coefficients of the polynomial

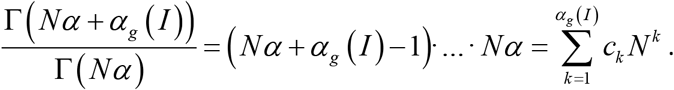

As mentioned earlier, we assume that the subjects’ estimate on each trial, 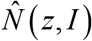, is given by the mean of the posterior distribution:

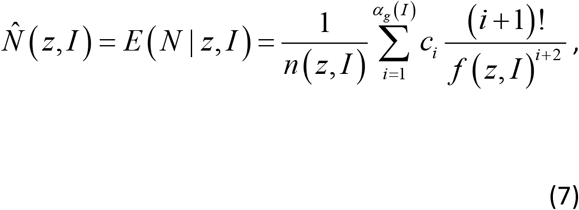

To determine whether this estimate follows Weber’s law, we again compute the variance of the estimate over trials conditioned on the true numerosity *N*. This expression is quite complicated to evaluate explicitly in general, but it simplifies considerably in the limit of large *N* (or equivalently large *z*). To see this, note that the mean of the likelihood (Eq. (1)) is given by

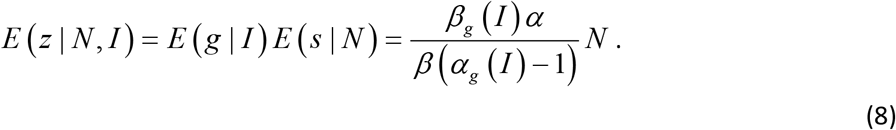

Therefore, in the limit of large *N, z* becomes large as well, *f* (*z*) becomes small and only the highest power of *f* (*z*) survives in Eq. (7). This implies that the posterior mean is given by:

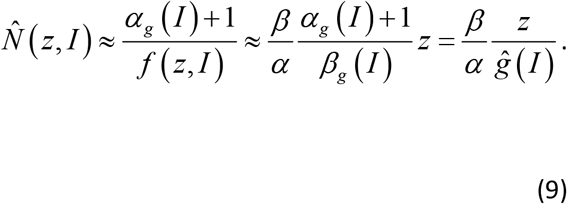

Where *ĝ* (*I*) is the maximum a posteriori estimate of *g* given *I* (*β*_*g*_ / (*α*_*g*_ +1) is indeed the mode of the inverse Gamma distribution). Again, 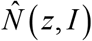 in Eq. (9) can be written as the product of two independent random variables 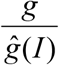 and 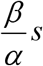, and its variance is similarly given by

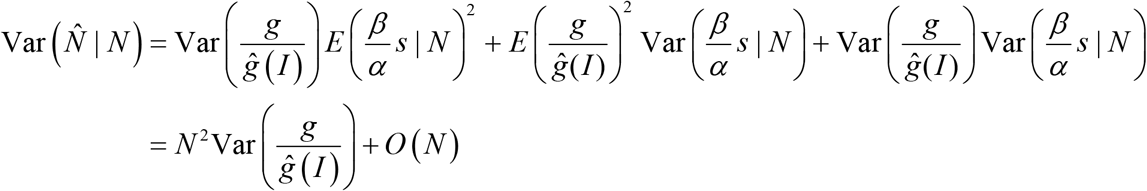

Therefore, for large *N*, this estimate follows Weber’s law, i.e., its variance is proportional to its mean squared, with a Weber fraction 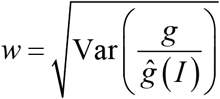, just as we observed with *α* = 1. Note also that in both of these cases, Weber’s law is caused by the fluctuation of the contrast estimate *ĝ* (*I*) around the true contrast *g*. This predicts that Weber’s law is still valid if the true contrast *g* is held fixed so long as the contrast estimate *ĝ* (*I*) is fluctuating.

#### Gamma Distributed s

The same line of reasoning that applies to counting discrete objects also applies to estimate of continuous quantities. In particular, if we assume a standard scale mixture model in which the quantity of interest *s* is a positive valued random variable and our observation, *z*, is a scaled version of *s*.

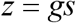

Here, *s* can be thought of as a feature of a natural image which can be linearly extracted from pixels and *g* plays the role of contrast. In the interest of analytic tractability we will assume that *s* is Gamma distributed and *g* is inverse Gamma, i.e.

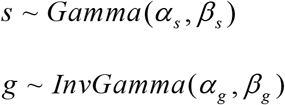

Note that we have suppressed the *I* dependence of *α*_*g*_ and *β*_*g*_ in the interest of clarity. Regardless, when this is the case, it is easy to show that the observation is distributed according to

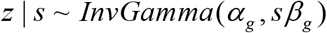

and

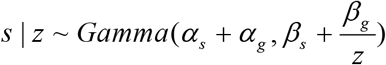

For Gamma distributed random variables, the ratio of the standard deviation to the mean is the square root of the inverse of the *α* parameter, i.e. (*α*_*s*_ + *α*_*g*_)^−1/2^. Since this parameter is independent of the observation, *z*, this ratio is constant and Weber’s law results when this generative model is correct. As in the previous case, we can also show that Weber’s law results when the generative model is incorrect but still used by the subject for inference. To see this more clearly, we again assume that the image contains a great deal of information about the contrast nuisance parameter *g* and both *α*_*g*_ and *β*_*g*_ are large. The posterior mean estimate of the quantity of interest *s* is then given by

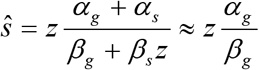

Note also, that once again, the estimate of *s* is obtained by normalizing *z* by the estimate of the gain or contrast 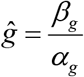. Conditioned on *s* the expectation and variance of this estimate are

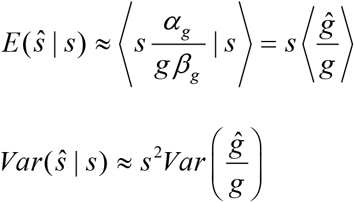

and so Weber’s Law behavior is observed.

#### Gaussian Distributed s

Precisely the same line of reasoning applies to Gaussian scale mixture models where the underlying quantity of interest, *s*, is a unit Gaussian random variable scaled by the square root of a Gamma distributed random variable random variables, i.e.

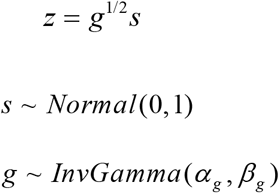

In this case, we can take advantage of the fact that

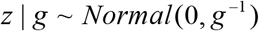

to conclude that

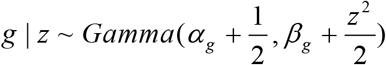

And therefore,

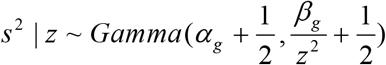

As in the previous case, a posterior of this form can be used to show that the ratio of the standard deviation to the mean of the posterior on *s*^2^ or absolute value of *s* does not depend on z, so that once again Weber’s Law results. In fact, one can show that estimates of |*s*|^*p*^ also obey Weber’s law for any *p* greater than or equal to 1. Regardless, we can once again consider estimates of *s* by taking the mean of the posterior, which in this case takes the form,

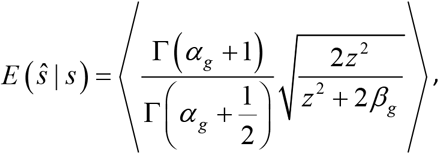

where the expectation is over *z* and the *α*_*g*_ and *β*_*g*_ pairs. Once again, a case of particular interest is when we have a lot of information about the contrast nuisance parameter, *g*, which implies that *α*_*g*_ and *β*_*g*_ are both large. When this is the case it is again easy to see that estimate of the magnitude of *s* is obtained is obtained by normalizing the magnitude of the observation by the an estimate of the contrast *g*^1/2^, that is

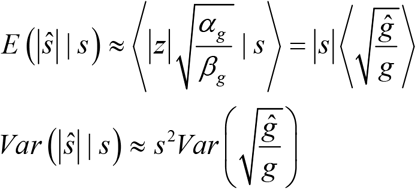

and so Weber’s Law holds once again.

### 2) Weber’s law due to learning a categorization bound

Another nuisance parameter that can give rise to Weber’s law is a categorization bound separating objects to be counted over from objects to be discarded. In a naturalistic setting this decision bound has to be inferred from sensory experience and fluctuations in the categorization bound will induce correlations among the individual acceptance probabilities of each object.

As an example, consider the problem of counting dogs in a display in which a variety of intermingled dogs and cats are shown. For each animal (indexed by *i*), we will assume that the visual system extracts a vector of sensory features, **x**, which follows a normal distribution with mean 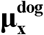 and 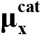 mean and identical covariance matrix Σ. In this situation, the optimal classifier is linear in the features and the subject needs to perform a classification of the form, *y*_*i*_ = **w**·**x**_*i*_ > *δ*, where **w** and *δ* are parameters to be learned.

Counting the number of dogs, *n*_*d*_, only requires to sum the number of animals for which *y*_*i*_ > *δ*. Next, we show that if **w** and *δ* are fixed and the fluctuations in **x**_*i*_ are independent across objects, the variance of the optimal estimate of *n*_*d*_ grows proportional to *N* which is to say that it does not follow Weber’s law. By contrast, if **w** or*δ* vary from trial to trial due for instance to online learning, this creates additional variability in *δ* or *y*_*i*_, in which case the estimate of *n*_*d*_ follows Weber’s law.

To see this, consider individual probabilities of the animals to be classified either correctly or incorrectly. Let us assume that the observation noise *p* (**x**_*i*_ | **μ**) is independent across animals. The probability *p*_11_ for a dog to be correctly classified is given by

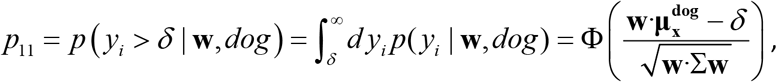

where Φ(*z*) is the cumulative normal distribution. Similarly the probability classified as a dog is given by *p*_10_ for a cat to be incorrectly

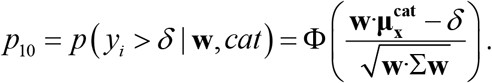

The expectation value and variance of the estimated number 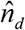 given the true number *n*_*d*_ and fixed **w** and *δ* will be

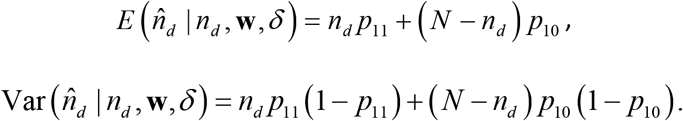

This confirms that for fixed **w** and *δ* the variance grows linearly with *N* which corresponds to counting noise but not Weber’s law. If however **w** and *δ* fluctuate from trial to trial, then *p*_11_ and *p*_10_ vary as well since they are functions of **w** and *δ*, which leads to:

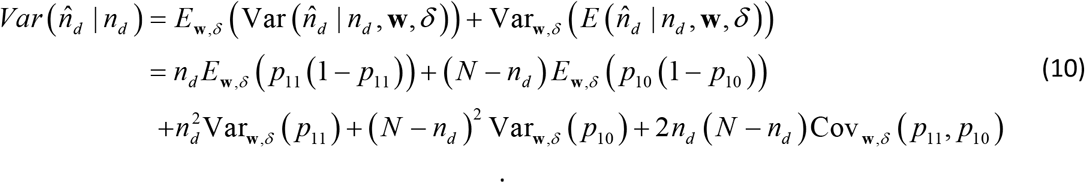

Additionally to the counting noise contribution in line 2 we get the contributions in line 3 due to the fluctuations in **w** and *δ*. The 3rd line grows proportionally to *N* ^2^ (i.e. Weber’s law is valid for large *N)* as long as the terms Var_**w**,*δ*_ (*p*_11_), Var_**w**,*δ*_ (*p*_10_) and Cov_**w**,*δ*_ (*p*_11_, *p*_10_) are positive and constant in *N*.

### 3. Efficient coding with input information following Weber’s law

In this section we illustrate how log-normal tuning curves provide the optimal encoding of a stimulus if the input to the neural population is already corrupted by Weber-law noise. To measure optimality, we minimize linear Fisher information, which corresponds to the variance of the locally optimal linear estimator (LOLE) in a discrimination task [1, 2].

Consider a population of independent Poisson neurons with Gaussian tuning curves tuned to a one-dimensional input *t*. If *t* is noisy as well, this will give rise to a population hill of activity shifting from trial to trial and information-limiting differential correlations [3].A Weber-like input noise can be achieved by the relation

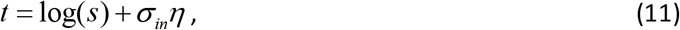

where *σ*_*in*_ corresponds to the Weber fraction and *η* is normally distributed.

Conditioned on *t*, we assume that the neural responses **r** are distributed according to

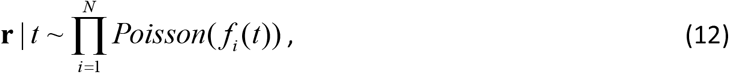

where the tuning curves are given by

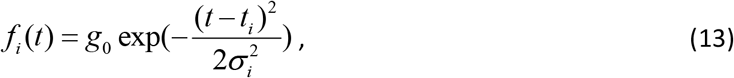

where *g*_0_ corresponds to the gain, *t*_*i*_ to the preferred input and *σ*_*i*_ to the width of the tuning curve. Note that different neurons do not have the same tuning curve width; we instead allow the tuning curve width to vary slowly according to a smooth function *σ*_*i*_ = *σ* (*t*_*i*_).

Since the equations (11) and (12) describe a doubly stochastic process, we can calculate the tuning curve and covariance matrix conditioned on the stimulus *s*. The tuning curves conditioned on *s* are given by:

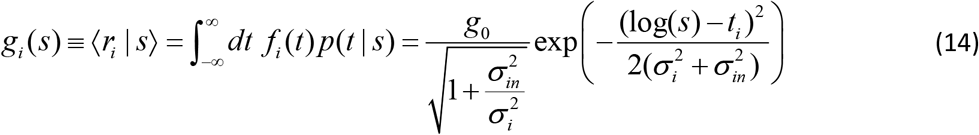

The covariance matrix is given by

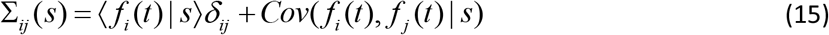

The first term is equal to *diag*(*g*_*i*_ (*s*)), similar to equation (14). The second term can be calculated analytically, but the resulting expression for Σ_*ij*_ (*s*) is difficult to invert. Instead we approximate (see justification in the next subsection)

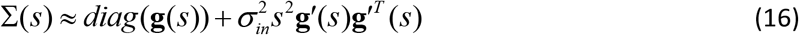

Using the Sherman-Morrison formula, we can calculate the linear Fisher information:

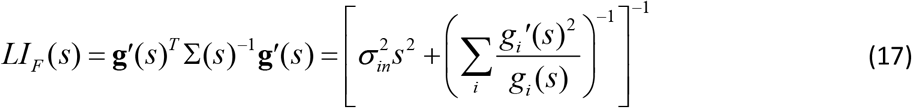

The first term in the curly brackets is caused by the input noise and the second term by the internal Poisson noise. By approximating the sum by a continuous integral we can further simplify the Poisson term:

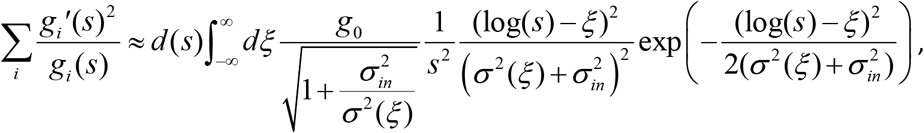

where *d*(*s*) is the density of neurons with preferred tuning log(*s*). This integral becomes tractable if we approximate the width function *σ* ^2^ (*ξ*) by a constant *σ* ^2^ (log(*s*)), an approximation which is valid if the width function changes slowly [4]. We obtain

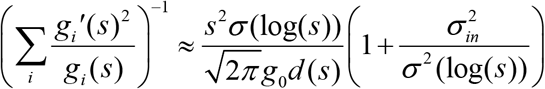

and

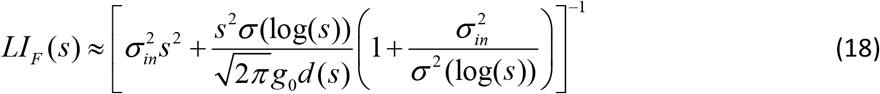

To find the optimal tuning curve width we maximize eq. (18) with respect to *σ* (log(*s*)) point-wise at each stimulus value *s* and find that the optimal width function is constant. The optimal value is given by:

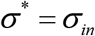

Thus, the linear Fisher information in the population is maximized if the width of the tuning curves is constant in log-space, as is the case in log-normal tuning curves. Note that since the optimization of (18) can be done separately for each stimulus value *s*, our argument does not rely on a specific form of a prior stimulus distribution.

We can go further and also compute the optimal density *d* (*s*) in the population, under a constraint on the number of neurons ∫ *d* (log(*s*)) ds and a constraint on the number of evoked spikes: *g*∫ *d* (log(*s*))*σ* (log(*s*)) p(s), where p is the prior density of the stimulus. If we optimize the objective function: ∫ log(*LI*_*F*_ (*s*)) *p*(*s*) ([4]), the optimal density is found after some calculations to be:

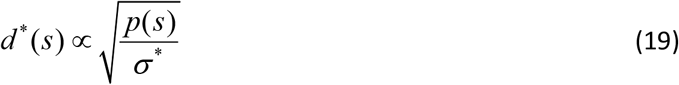

#### Justification for the approximation in eq. (16)

In order to justify the approximation of the covariance matrix in eq. (16), we consider for a moment an alternative population where all neurons are deterministic instead of being Poisson. The covariance matrix in this second population is *Cov*(*f*_*i*_ (*t*), *f* _*j*_ (*t*) | *s*), but, because of the data-processing inequality, the Fisher information in that noiseless population would still be limited by the information in *t*. Applying the Cramer-Rao inequality to the vector function (*f*_1_(*t*),…, *f*_*n*_ (*t*)), we can derive a lower bound on the covariance of the noiseless population:

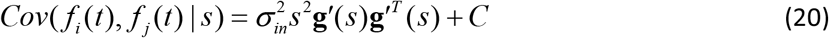

where *C* is an unknown positive semi-definite matrix.

Therefore, the covariance matrix of the Poisson population can be expressed as:

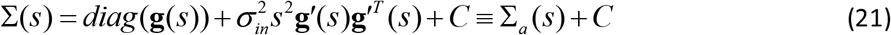

Applying the Woodbury identity twice and denoting Σ_*a*_ (*s*) the approximation in the left hand side of eq. (16), the inverse is given by:

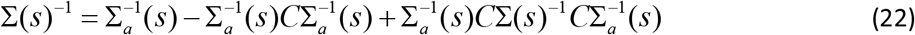

Consider the vector V with elements: 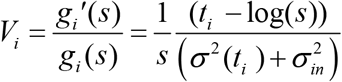. We now prove that *V*^*T*^ *CV* ≈ 0, which is equivalent to *CV* ≈ 0, since *C* is positive semi-definite. This implies that the second and third term in eq. (22) do not contribute to the full linear Fisher information **g**′(*s*)^*T*^ Σ(*s*)^−1^**g**′(*s*), since 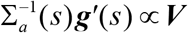. Finally, the linear Fisher information is

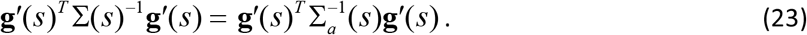

This identity justifies the use of our approximation in eq. (16).

To prove that *V*^*T*^*CV* ≈ 0, we will compute the variance of the variable ***V***^*T*^ ***f*** (*t*) conditioned on s in two different ways. On one hand:

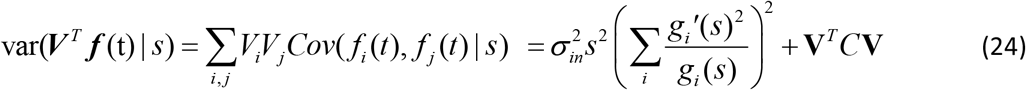

On the other hand, we can compute ***V***^*T*^ ***f*** (*t*) by approximating the sum by a continuous integral:

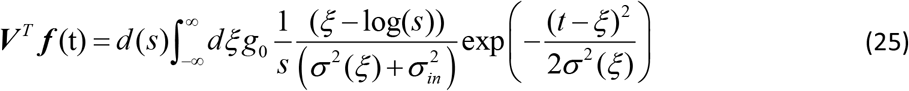

Approximating again that: *σ* ^2^ (*ξ*) ≈*σ* ^2^ (log(s)), this integral becomes tractable and we find that:

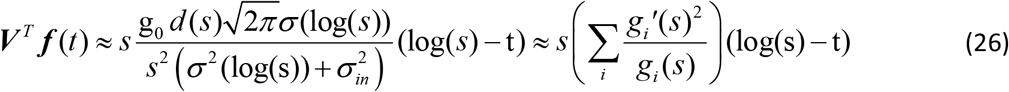

Therefore, the variance is given by

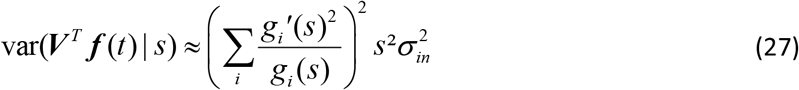

Equating eq. (24) and (27) we obtain:

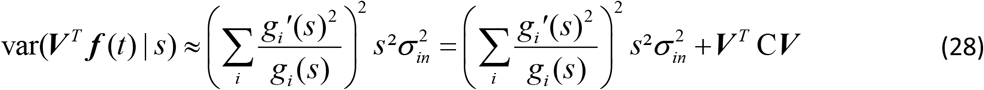

From this we conclude that indeed ***V*** ^*T*^ C***V*** ≈ 0, which justifies the use of the approximation.

